# Do count-based differential expression methods perform poorly when genes are expressed in only one condition?

**DOI:** 10.1101/017673

**Authors:** Xiaobei Zhou, Mark D. Robinson

**Affiliations:** Institute of Molecular Life Sciences, University of Zurich, CH-8057 Zurich,Switzerland; SIB Swiss Institute of Bioinformatics, University of Zurich, CH-8057 Zurich,Switzerland

## Abstract

A Correspondence to **Comprehensive evaluation of differential gene expression analysis methods for RNA-seq data** by Rapaport F, Khanin R, Liang Y, Pirun M, Krek A, Zumbo P, Mason CE, Socci ND and Betel D, Genome Biol 2013, 14:R95

## Background

Statistical methods for determining transcriptional changes between (replicated) groups of cell populations using RNA sequencing (RNA-seq) data are now quite mature. Several themes that emerged from the past decade of modeling microarray data apply analogously to RNA-seq data: parameter moderation is critical, multiple testing corrections are necessary and flexible frameworks (e.g. linear models) to account for the effect of covariates are essential. In RNA-seq data, popular packages such as edgeR, DESeq and DESeq2 [8, 1, 5] perform detailed modeling of the dispersion-mean relationship, with variations on fitting a dispersion by mean trend and moderating estimates toward the trend. Likewise, careful modeling of the mean-variance relationship of transformed data has been proven effective, essentially “unlocking” the world of heteroskedastic linear regression [3].

A recent report in Genome Biology from Rapaport and co-authors claimed that some methods, namely PoissonSeq [4] and limma [9], “have improved modeling of genes expressed in one condition”, where they showed a striking difference in the ability to separate differential expression [7]. From a methodological perspective, this result caught our interest and prompted us to understand how aspects of all-zero-in-one-condition manifest undesirable properties in count-based models. Briefly, we: i) found that Rapaport et al. made an error in the signal-to-noise (S/N) calculation for edgeR; and, ii) a re-analysis suggests that count-based methods perform as well or better than other methods, counter to the original conclusion.

The Rapaport manuscript is an excellent model of modern bioinformatics research, in terms of making processed data and code available that reproduce figures from their manuscript. In many cases, the small details can be important and this open source model facilitates quick access to understand precisely what settings were used. We fully support this model and by default, also make our code available. In this Correspondence, we investigate the genesis of differences in method performance that Rapaport and co-authors observed and provide our view of how performance results can be sensitive to decisions made.

## Genes expressed in only one condition

We first briefly summarize the analysis that Rapaport and colleagues reported, with respect to the all-zero-in-one-condition case.

Using gene-level read counts, they isolated genes that exhibit zero counts across all replicates of a single condition; in general, the number of such genes is related to the depth of sequencing dedicated to each sample, with deeper sequencing resulting in fewer such cases. The dataset in question, comparing GM12892 cells to H1-hESC cells [13], with 3 and 4 replicates, respectively, had typical read depths (16-39 million mapped reads) for such experiments. They used the following pipeline: i) from the count table, generate DE P-values for several methods; ii) calculate S/N using “normalized” data; iii) plot negative log P-value versus S/N, where they expect a monotonic positive dependency (correlation); and, iv) generate receiver operating characteristic (ROC) curves with thresholds on the S/N to illustrate the ability to separate low S/N (<3) from high S/N (>3).

They highlighted that count-based methods such as DESeq and edgeR, which infer changes in expression via the negative binomial (NB) model, do not perform very well in this case. It is worth noting that this is a non-standard use of ROC curves: here, all genes are strictly DE, but they vary in their magnitude of change. So, the ROC curve represents the ability to separate low S/N from high S/N. Rapaport and colleagues postulated that the NB model reduces to Poisson (dispersion *≈* 0) and lacks the ability to handle the “wide variations” in gene counts among replicate libraries. Our aim with this report is to understand the origins of this result, whether it be a shortcoming of the dispersion estimation strategy or in the inference machinery since parameter estimates are on the boundary of the parameter space.

## Signal-to-noise has some potential limitations

We became interested in the suitability and robustness of the S/N metric itself, since it forms the basis for the “truth” in Rapaport’s ROC result. In theory, the S/N of the non-zero observations should accurately reflect the significance of model-based P-values for the expressed-in-one-condition versus zero differences. In practice, however, there are some potential difficulties: the sample sizes are small and therefore, the S/N itself is subject to considerable estimation uncertainty; it is well known in count data that the variance is intimately tied to the mean, so it is not clear whether S/N should be calculated on the linear scale. In addition, a notable aspect of the Rapaport ROC comparison is that while the same S/N cutoff (=3) is used across all methods, different sets of true and false DE labels are used; this makes the curves difficult to compare, since both the truth and score change by method. We explore these issues here.

Table 1 and Figure 1 give illustrative examples of the differences in the originally-calculated S/N between edgeR and voom. Figure 1 gives a scatter plot of S/N calculated on each method’s normalized data, highlighting in some cases large differences. Table 1 shows the top 10 genes each, in terms of edgeR’s (estimated) false discovery rate (FDR) and calculated S/N, respectively (The full table of zero counts, differential statistics and S/N is given in Additional file 1). Here, it is evident that several genes that show little evidence for differential expression, have very high S/N for edgeR but not for voom (e.g. C17orf66, TM4SF19, NPY1R). However, the P-values seem to appropriately reflect the magnitude of evidence for DE, although they are on drastically different scales between edgeR and voom (see Discussion for further commentary on this). In addition, several genes that show the largest evidence against the null hypothesis (e.g., PLEK, MS4A1, etc.) show relatively low S/N for edgeR and would be counted as false discoveries (according to a S/N=3 cutoff), while voom’s higher S/N would result in these counted as true positives. Therefore, it is not clear whether the ROC curve is reflecting the accuracy of the S/N calculation itself or of the statistical method’s capabilities. Upon investigation, the differences in S/N exhibited in Figure 1 resulted from a code error in the original report (See Additional file 2, Figure 1).

**Figure 1:**
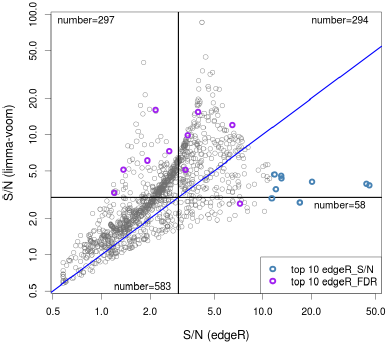
Scatter plot of S/N (signal-to-noise) for limma-voom and edgeR for the ENCODE dataset, where S/N is calculated from the non-all-zero condition. More information for the colored points is given in Table 1.

**Table 1:** A table of selected genes according to the top 10 originally-calculated S/N (for edgeR-normalized data; first 10 rows) and top 10 FDR for differential expression (edgeR P-values; second 10 rows). The table includes the count-per-million table (A=GM12892, B=H1-hESC), S/N and estimated false discovery rate (FDR) for edgeR and limma-voom for the ENCODE dataset comparing 3 replicates of GM12892 to 4 replicates of H1-hESC.

| id | A1 | A2 | A3 | B1 | B2 | B3 | B4 | edgeR S/N | edgeR FDR | voom S/N | voom FDR |
| --- | --- | --- | --- | --- | --- | --- | --- | --- | --- | --- | --- |
| MIPOL1 | 0.0 | 0.0 | 0.0 | 237.1 | 232.5 | 226.0 | 227.5 | 45.75 | 7.47e-31 | 3.79 | 3.32e-07 |
| AQP4 | 0.0 | 0.0 | 0.0 | 46.1 | 45.0 | 46.7 | 44.4 | 43.87 | 1.44e-14 | 3.90 | 1.99e-06 |
| FAM19A4 | 0.0 | 0.0 | 0.0 | 142.1 | 131.1 | 143.8 | 131.4 | 20.21 | 1.91e-24 | 4.06 | 4.72e-07 |
| C17orf66 | 2.3 | 2.1 | 2.1 | 0.0 | 0.0 | 0.0 | 0.0 | 16.96 | 1.89e-02 | 2.72 | 7.40e-04 |
| TM4SF19 | 3.5 | 3.2 | 3.2 | 0.0 | 0.0 | 0.0 | 0.0 | 16.96 | 3.00e-03 | 2.72 | 2.39e-04 |
| SOX1 | 0.0 | 0.0 | 0.0 | 7.5 | 6.7 | 6.5 | 6.3 | 13.02 | 8.37e-05 | 4.33 | 1.27e-04 |
| HPGD | 0.0 | 0.0 | 0.0 | 22.6 | 21.0 | 19.6 | 19.0 | 12.97 | 5.08e-09 | 4.56 | 7.48e-06 |
| LOC100131176 | 0.0 | 0.0 | 0.0 | 17.9 | 15.3 | 18.7 | 17.2 | 12.02 | 5.14e-08 | 3.51 | 1.49e-05 |
| ZNF385D | 0.0 | 0.0 | 0.0 | 135.5 | 155.0 | 155.0 | 132.3 | 11.80 | 2.60e-25 | 4.67 | 3.70e-07 |
| NPY1R | 0.0 | 0.0 | 0.0 | 209.8 | 179.9 | 179.3 | 208.5 | 11.38 | 1.33e-27 | 2.95 | 6.77e-07 |
| PLEK | 25082.8 | 12622.5 | 11394.8 | 0.0 | 0.0 | 0.0 | 0.0 | 2.16 | 1.79e-216 | 16.11 | 9.36e-09 |
| MS4A1 | 25455.1 | 14937.7 | 12886.8 | 0.0 | 0.0 | 0.0 | 0.0 | 2.63 | 2.62e-215 | 7.26 | 1.60e-08 |
| SLAMF1 | 7407.2 | 4859.3 | 4283.2 | 0.0 | 0.0 | 0.0 | 0.0 | 3.32 | 2.98e-165 | 5.11 | 2.53e-08 |
| CCL3 | 11057.5 | 3413.1 | 3544.1 | 0.0 | 0.0 | 0.0 | 0.0 | 1.37 | 4.62e-165 | 5.13 | 2.15e-08 |
| FCRLA | 7742.0 | 2979.1 | 3879.8 | 0.0 | 0.0 | 0.0 | 0.0 | 1.92 | 1.01e-161 | 6.08 | 1.84e-08 |
| RGS1 | 9939.5 | 9967.3 | 7741.6 | 0.0 | 0.0 | 0.0 | 0.0 | 7.22 | 4.53e-159 | 2.66 | 1.44e-07 |
| DPPA4 | 0.0 | 0.0 | 0.0 | 14580.2 | 15215.1 | 14745.3 | 10617.3 | 6.47 | 2.37e-158 | 12.02 | 1.84e-08 |
| TDGF1 | 0.0 | 0.0 | 0.0 | 15699.8 | 15481.1 | 13374.5 | 8522.5 | 3.98 | 6.37e-157 | 15.48 | 1.84e-08 |
| SFRP2 | 0.0 | 0.0 | 0.0 | 14673.3 | 15229.5 | 13067.2 | 7234.4 | 3.43 | 1.84e-153 | 9.87 | 2.15e-08 |
| BLK | 9943.0 | 2954.7 | 2351.8 | 0.0 | 0.0 | 0.0 | 0.0 | 1.20 | 2.98e-147 | 3.28 | 5.17e-08 |

Another aspect to understand is the scale on which the S/N is calculated on. As is well known with count data, the variance is related to the mean. In particular, using the NB parameterization with mean *µ* and variance *µ*(1 + *µ* • *φ*), the theoretical S/N is then:

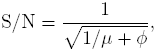

which implies 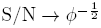 with sufficiently large *µ*. Thus, depending on the mean, the S/N calculation is capturing (inverse square root of) dispersion. For the ENCODE data, this relationship is shown in Additional file 2, Figure 2. Since the S/N calculations are most relevant when the variance is independent of the mean, we explored how transforming the data, which alters the mean-variance relationship, affects the results of the ROC comparisons that Rapaport and co-authors performed. Figure 2 (a)-(c) show mean-variance relationships for S/N calculated on different scales and (d)-(f) highlight their corresponding ROC performances. In all cases, the true/false labels for the ROC curves are the same across methods; specifically, counts-per-million from edgeR are used to base the S/N calculation. Since the scale of data changes the scale of S/N, true genes are selected according to S/N *>* 40^*th*^ percentile and false as the lowest 20% of S/N to give a grey zone of uncertainty in the middle (Additional file 2, Figure 3 gives alternative settings for these cutoffs, but results are unaffected). Figure 2 (d) shows similar results to the original Rapaport study, whereas Figure 2 (e),(f) show a remarkable reversal in performance, giving clear evidence for our earlier concern regarding the S/N calculation.

**Figure 2:**
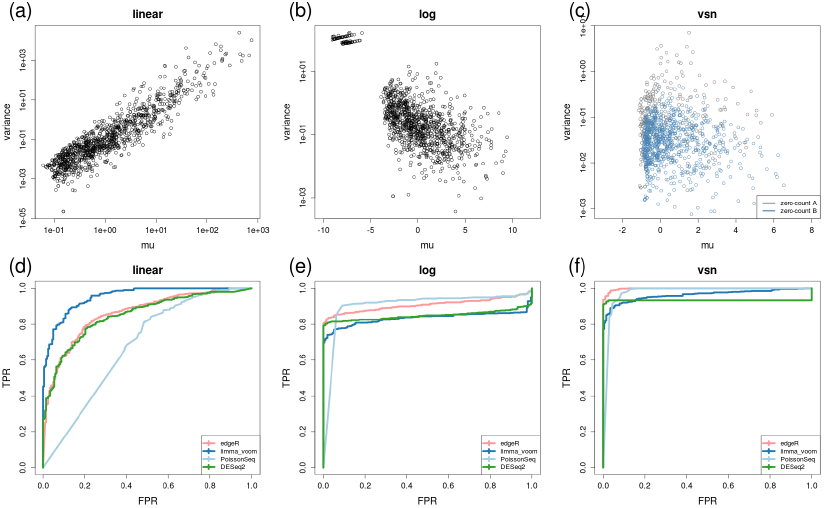
Panels (a)-(c) give mean-variance relationships for different scales of the original all-zero-in-one-condition data. Panels (d)-(f) give corresponding ROC curves for the ENCODE dataset (GM12892 cells to H1-hESC), using S/N to set the true labels. Here, the signal-to-noise (S/N) is calculated from (TMM-normalized) counts-per-million and used for all methods. “linear” is equivalent to Rapaport’s method, where S/N is calculated on the counts-per-millions. “log” represents S/N calculated on log-transformed counts-per-million. “vsn” represents S/N calculated on variance-stabilized data([2]). ROC curves employ the same labels across all methods: the top 40% of S/N are used as true DE genes whereas the lowest 20% as false. Each method’s P-value is used for ranking the genes.

## Count-based methods perform well on zero-in-one-condition simulation

Given recent efforts in simulating RNA-seq count table [11, 10, 12], we tried to create a representative simulation for the zero-in-one-condition situation. The simulation was designed as follows: i) generate a dataset with no DE, ii) randomly select genes across the spectrum of expression levels and set counts for one condition (chosen at random) equal to zero to represent “true” DE genes. As previously, we sampled NB mean and dispersion estimates from the joint distribution of estimates using a large dataset (here, from [6]) and filtered out extreme dispersion values. Altogether, 30, 000 features were generated, in a 5 versus 5 two-group comparison and zero counts were introduced to 5% of the features. To reflect the fact that zeros occur somewhat more often at lower expression across various datasets (See Additional file 2, Figure 4), we increased the frequency of zero-counts at low expression value.

Based on the results of this simulation (Figure 3), ROC curves with the method’s 5% FDR highlighted (panel a) and plots of true positive rate versus achieved FDR (panel b), we again see that count-based models perform well in the zero-in-one-condition situation. In addition, we explored the postulation that the NB model gets reduced to Poisson in these zero-count situations. By comparing the dispersion estimates calculated from the single non-zero condition to the original non-zero-in-both conditions data, it does not appear that the dispersion estimates are drastically reduced (see Additional file 2, Figure 5).

**Figure 3:**
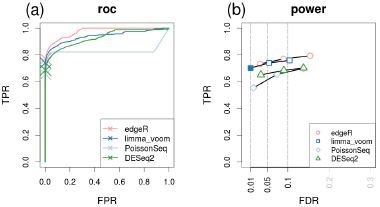
(a) ROC curves and (b) true positive rate (TPR) versus “achieved” FDR curves of DE methods for the simulation dataset with zero-counts introduced as the true DE genes (overall performance of 3 simulations); achieved FDR is the actual rate of false discoveries at the corresponding cutoff and this rate should ideally be controlled at the desired level. For the ROC curves, the X on each curve represents the method’s TPR at the (estimated) 5% FDR cutoff. For the TPR versus achieved FDR curves, points are plotted at the following cutoffs: 0.01, 0.05 and 0.1; filled-in points represent that the method has correctly controlled the error rate at the cutoff.

## Discussion

As developers and users of bioinformatics strategies, we are particularly interested in the metrics and methods that differentiate performance between the available tools. In this paper, we claim that count-based methods perform well in the situation where genes are only expressed in one condition, in contrast to an earlier report. We showed that a code error and the chosen scale of S/N resulted in the earlier conclusion that count-based methods suffer performance in this situation. By calculating the S/N on a different scale and using the same set of labels across methods, a reversal of method performance was observed. This highlights a sensitivity to decisions made in constructing the benchmark.

Using a customized simulation that introduces zero-counts in one experimental condition, we demonstrated that the performance of count-based is actually on par or better than other methods. We also debunked the postulation that poor performance is related to dispersion estimation in count models.

In the process of seeking out the origins of this statistical performance difference, we discovered another potentially interesting phenomenon that may affect the interpretation of results. Looking at Table 1 and Additional file 1, it is evident that the scale of P-values is drastically different between edgeR and voom. Although this observation appears rather unrelated to the ability to separate true from false DE genes, it is an indication that the scale of observations modeled affects the magnitude of statistical evidence derived. Not surprisingly, method performance is ultimately dependent on the scales, parameters, datasets used for the evaluation.

## Supplemental material

Additional Supplementary material is available from: http://imlspenticton.uzh.ch/robinson_lab/zero_count/

## Author’s contributions

MDR and XZ both designed and conducted analyses and wrote the manuscript.

## Acknowledgements

X.Z. is supported by SNSF project grant (143883). M.D.R. acknowledges funding from the European Commission through the 7th Framework Collaborative Project RADIANT (Grant Agreement Number: 305626). We acknowledge data processing help from Malgorzata Nowicka at the outset of the project and to her and Charity Law for helpful feedback on an earlier version of the manuscript.

## Tables

### Additional Files

#### Additional file 1 — Table of statistics for zero-count genes

table of zero counts, differential statistics and S/N for the ENCODE dataset

#### Additional file 2 — Supplementary Figures

This file contains the mentioned Supplementary Figures

#### Additional file 3 — R code and data

This file contains R code and necessary data used to reproduce the figures in the main manuscript and in the supplement.

